# Recording of “sonic attacks” on U.S. diplomats in Cuba spectrally matches the echoing call of a Caribbean cricket

**DOI:** 10.1101/510834

**Authors:** Alexander L. Stubbs, Fernando Montealegre-Z

## Abstract

Beginning in late 2016, diplomats posted to the United States embassy in Cuba began to experience unexplained health problems—including ear pain, tinnitus, vertigo, and cognitive difficulties^1–4^—which reportedly began after they heard^1,2^ strange noises in their homes or hotel rooms. In response, the U.S. government dramatically reduced^1–3^ the number of diplomats posted at the U.S. embassy in Havana. U.S. officials initially believed^1,2,5^ a sonic attack might be responsible for their ailments. The sound linked to these attacks, which has been described as a “high-pitched beam of sound”, was recorded by U.S. personnel in Cuba and released by the Associated Press (AP). Because these recordings are the only available non-medical evidence of the sonic attacks, much attention has focused on identifying health problems^6–11^ and the origin^12–17^ of the acoustic signal. As shown here, the calling song of the Indies short-tailed cricket (*Anurogryllus celerinictus)* matches, in nuanced detail, the AP recording in duration, pulse repetition rate, power spectrum, pulse rate stability, and oscillations per pulse. The AP recording also exhibits frequency decay in individual pulses, a distinct acoustic signature of cricket sound production. While the temporal pulse structure in the recording is unlike any natural insect source, when the cricket call is played on a loudspeaker and recorded indoors, the interaction of reflected sound pulses yields a sound virtually indistinguishable from the AP sample. This provides strong evidence that an echoing cricket call, rather than a sonic attack or other technological device, is responsible for the sound in the released recording. Although the causes of the health problems reported by embassy personnel are beyond the scope of this paper, our findings highlight the need for more rigorous research into the source of these ailments, including the potential psychogenic effects, as well as possible physiological explanations unrelated to sonic attacks.

---

Additional embassy personnel reported hearing sounds at night^1–5^ and many were sent to the U.S. for medical evaluation. A team from the University of Pennsylvania presented^4^ evidence of medical abnormalities. The U. Penn. paper has however been criticized as using an arbitrarily low threshold for neurological impairment^6–8^ and improperly ruling out potential causes such as functional neurological or psychological disorders^9–11^.

United States personnel made multiple recordings of the distinctive sound and these recordings were played to embassy personnel so they would know what to listen for^5^. The Associated Press (AP) received several of these recordings and posted^5^ one representative sample online. Recordings were sent^5^ to the U.S. Navy and FBI for analysis, and some were made available^12^ to the Cuban government. Because these recordings are the only non-medical evidence available on the “sonic health attacks” in Cuba, much attention has focused on identifying the origin of this acoustic signal, and on establishing whether it is connected to the reported health outcomes. A Cuban government report suggested^12^ that the Jamaican field cricket *Gryllus assimilis* was responsible. Other researchers posited that the noise might be^13^ the byproduct of a beam of high-power microwave-pulsed radiation. Another team suggested^14–17^ that intermodulation between ultrasound emitters could produce a spectral shape similar to the AP recording, and that this audio signal may be a byproduct of malfunctioning eavesdropping equipment.

After listening to the AP recording A.L.S. was reminded of his experiences conducting fieldwork in the Caribbean. The recording sounded like an insect, yet the pulse structure of the AP-released file^5^ does not look like^17^ classic oscillograms presented in the biological literature on insect calls. If an insect were responsible for the sounds recorded by U.S. personnel in Cuba, this should be verifiable by quantitatively comparing recordings of calling insects to the AP sample.

Male crickets produced their calls by wing stridulation. One wing bears a vein with systematically-organized indentations and the other a scraper. The wings open and close, but only during the closing phase does the scraper strike consecutively each file tooth and produce a pulse of sustained oscillations amplified by specialized wing cells. The entire sequence of oscillation is known as a syllable. Therefore, a syllable is made of a number of oscillations that match^31^ the number of teeth struck in the file. The structure of a pulse is affected by^32^ the duration of muscular twitch and varying tooth spacing, which in conjunction with wing deceleration cause a commensurate reduction in the frequency of tooth-strikes towards the end of the pulse. Therefore, all cricket pulses in nature exhibit^33^ a gradual reduction in the instantaneous frequency as the pulse evolves.

The recording released by the AP^5^ has a number of measurable parameters. The power spectrum resembles a picket fence (Figure 1) with most of the power concentrated around 7 kHz. The picket fence of emission occurs at integer multiples of the pulse repetition rate (PRR) of ∼180 Hz. The sound is continuous for the duration of the recording.

**Fig. 1.**
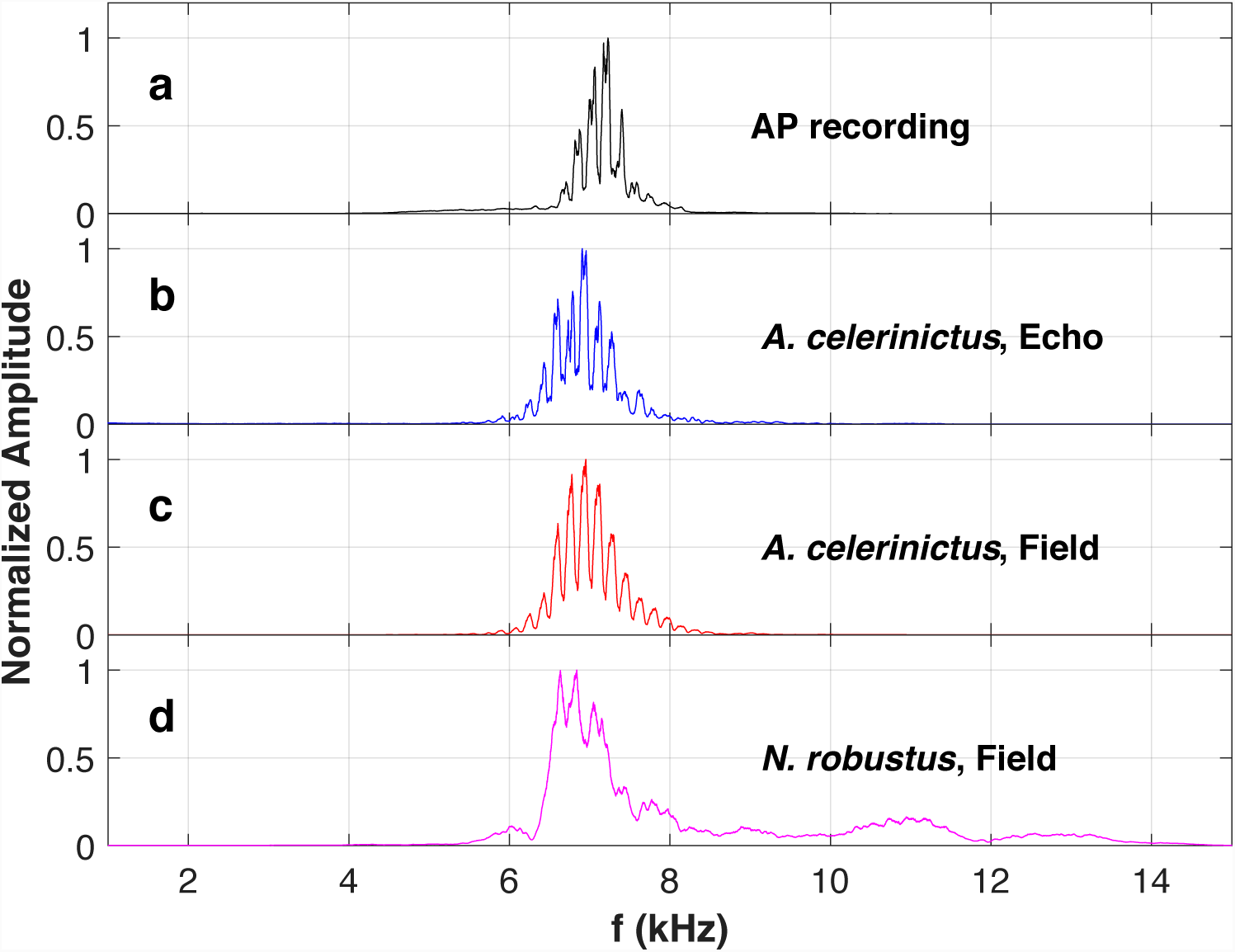
Normalized amplitude spectrum of potential sound sources compared to AP recording. **a**, the AP-released recording from Cuba, **b**, recording of *A. celerinictus* played on a speaker and recorded indoors, **c,** *A. celerinictus* and **d** the katydid *Neoconocephalus robustus* recorded in the field. *N. robustus* has a broader spectral emission, with more power at higher frequencies, than the cricket *A. celerinictus.* The distinctive picket fence structure in the amplitude spectrum occurs at integer multiples of the pulse repetition rate (PRR), in this case multiple of 180 for the A.P. recording and *A. celerinictus* recording with and without echoes. The picket microstructure depends on PRR stability. Any proposed source must conform to both the peak emission at 7 kHz and have sufficient PRR stability to produce this characteristic picket fence structure in the amplitude spectrum.

The combination of a definitive carrier frequency (7 kHz) and PRR (180 Hz) allows for an assessment of potential calling insect sources, as seen in Figure 2. A number of insect species are capable of producing a 7 kHz carrier frequency, whereas a PRR as high as ∼180 Hz is rare in nocturnal insects that produce continuous calls. The PRR of many insects varies with temperature, but the peak carrier frequency remains^17–24^ comparatively stable. After an extensive evaluation of online recordings, the katydid *Neoconocephalus robustus* (Scudder 1862) and cricket *Anurogryllus celerinictus* (Walker 1973) calls were downloaded^22^ and analyzed for a number of spectral parameters, as both were potential matches in carrier frequency and PRR (see Methods). Both insects can call continuously, they share a peak carrier frequency of ∼7 kHz, and are capable of^20–22^ a PRR of 180 Hz or above. Furthermore, *A. celerinictus* has^20^ the fastest PRR of any continuously-calling cricket in the Caribbean or North America, and *N. robustus* is^22^ the loudest insect sound known from North America.

**Fig. 2.**
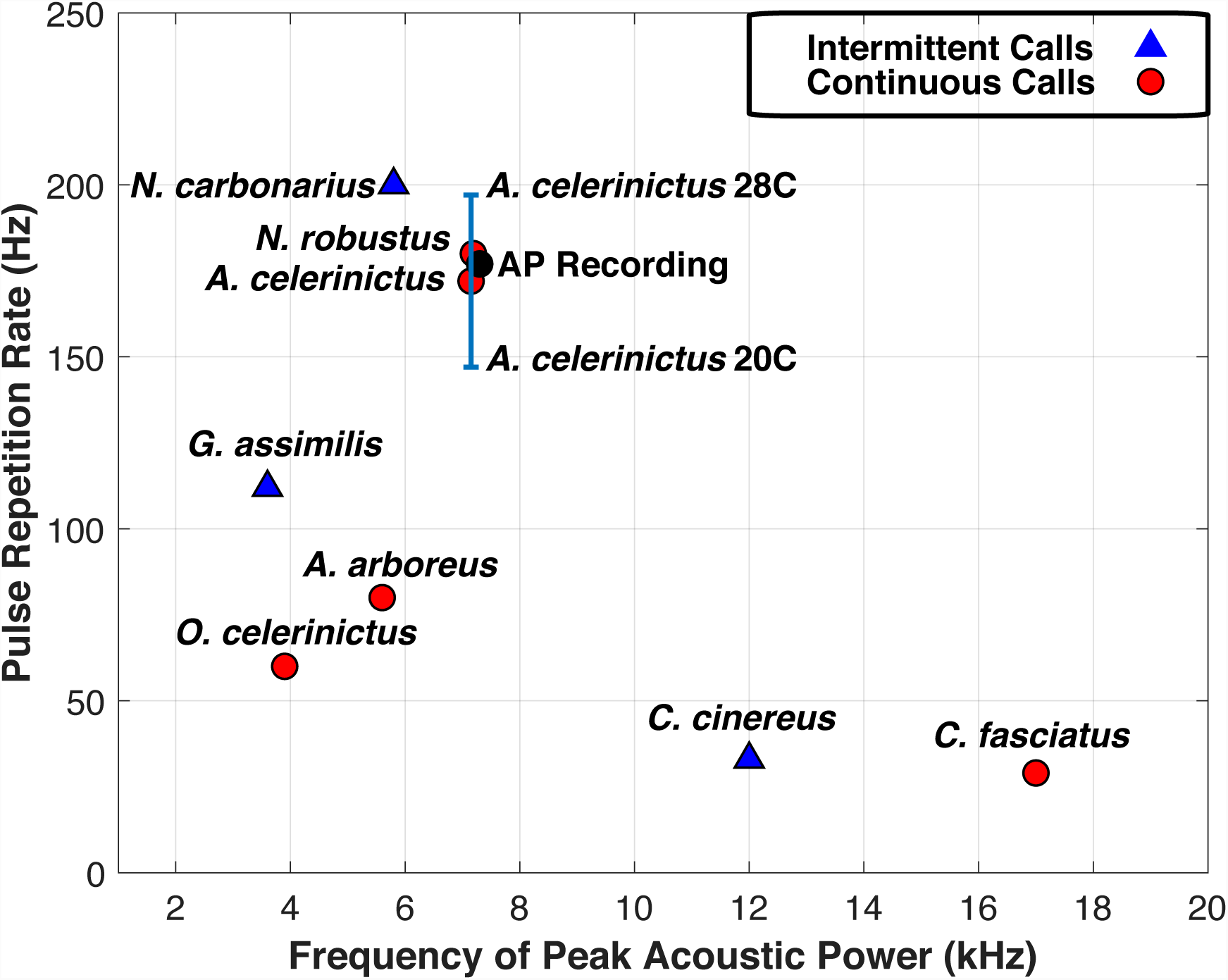
Pulse repetition rate vs. peak acoustic power frequency of various insect calls compared to the AP-released recording from Cuba. Both *A. celerinictus* and *N. robustus* are capable of continuously producing a sound with peak power at 7 kHz modulated at a PRR of ∼180 Hz. Continuously-calling insects are shown as circles while those with intermittent pulse trains are shown as triangles. The PRR’s for these insects are temperature dependent, as shown by the bar for *A. celerinictus.*

The Cuban government was given access^12^ to multiple recordings by the U.S. government. The Cuban report proposed^12^ that the cricket *G. assimilis* was responsible for this sound. This insect does not call continuously, but rather produces^22–24^ an intermittent somewhat melodic chirp once per second. U.S. personnel on the other hand reported^1,2^ and recorded^5^ a continuous high-pitched buzzing. Additionally, *G. assimilis* calls use a much lower peak carrier frequency of 3.6 kHz and a PRR of less than 120 Hz^3,6^, while the AP recording has a carrier frequency of 7 kHz (as do the other audio samples analyzed^12^ in the Cuban report) and a PRR of almost 180 Hz as seen in Fig. Given that the specific organism identified^12^ in the Cuban report fails on all quantitative metrics to explain the sound recorded in Havana, and would sound qualitatively different even to non-experts, it is understandable that U.S. authorities met this explanation with skepticism.

The picket fence structure in the power spectrum is determined by the stability of the PRR. This is shown in Figure 3.

**Fig. 3.**
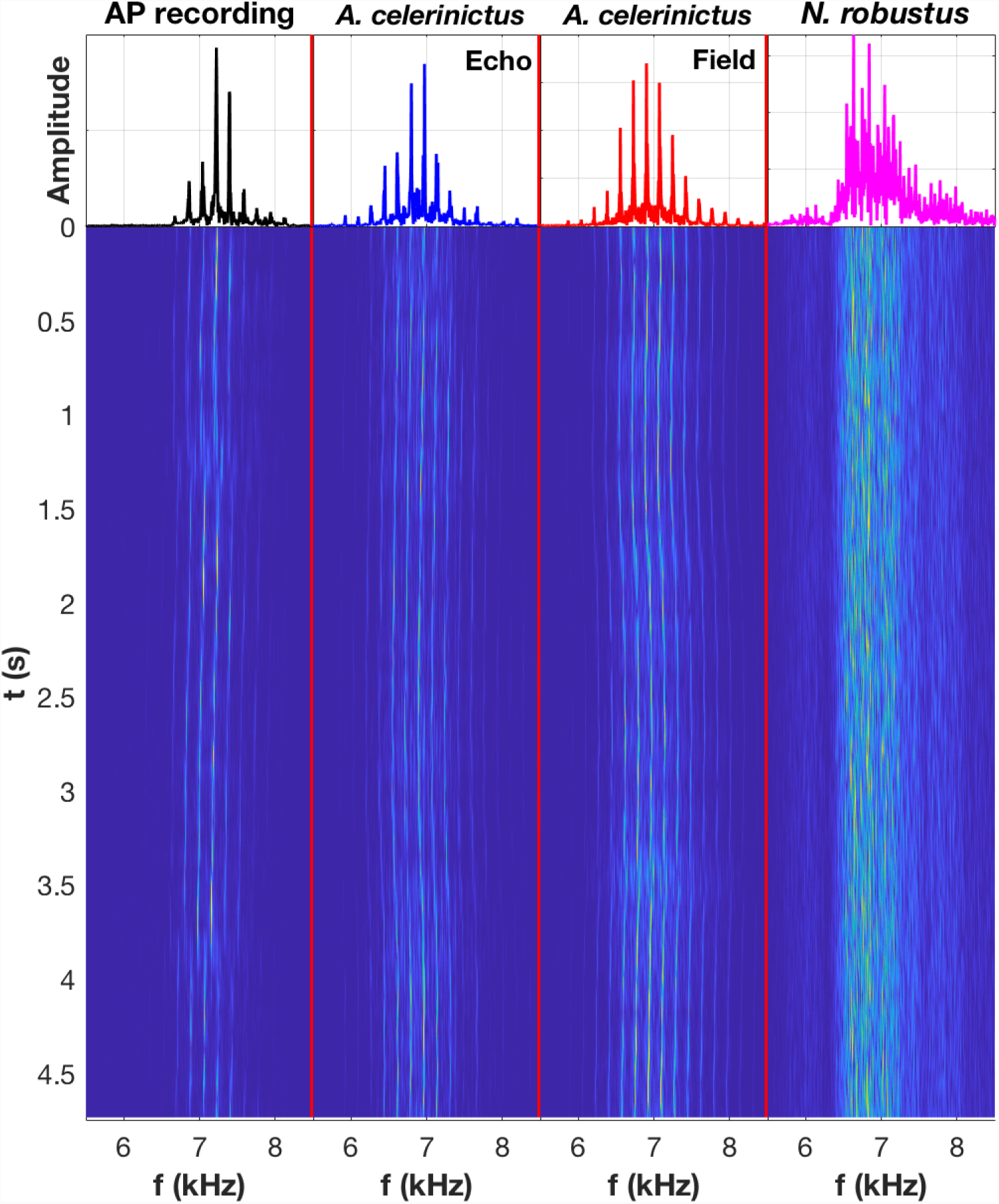
Plot of the pulse repetition rate stability. The lower panels show the evolution of the amplitude spectrum. Time runs vertically in the lower panel, and frequency increases to the right. The plots at the top show a cut across the waterfall diagram at t=0. Each line in the diagram comprises 8192 samples, spanning 186 msec. The AP recording and both recordings of *A. celerinictus* exhibit relatively (but not perfectly) stable PRR, whereas *N. robustus* shows significant short-term variability in PRR. The evolution of *A. celerinictus* matches the AP recording; both show few-percent fractional variations in PRR on a characteristic timescale of seconds.

The AP recording exhibits a non-uniform pulse structure (Figure 4) that at first glance is inconsistent with field and lab recordings^17–22^ of calling insects. The AP recording has sufficient variation in PRR (such as the offsets visible at 1.25 and 4 seconds) that it is unlikely to have been generated by a regulated digital signal source. U.S. personnel in Cuba reported hearing^1,2^ these sounds indoors. Ricocheting sound off walls, floors, and ceilings could produce complicated interference patterns or “echoes” obscuring the original pulse structure. To test if this might explain the pulse structure in the AP sample, a simple experiment was conducted. The *A. celerinictus* field recording was played on a high-fidelity loudspeaker, and recordings were made at various locations indoors. The pulse structure of a representative recording is shown in Fig. 4 and Extended Data Figure 1.

**Fig. 4.**
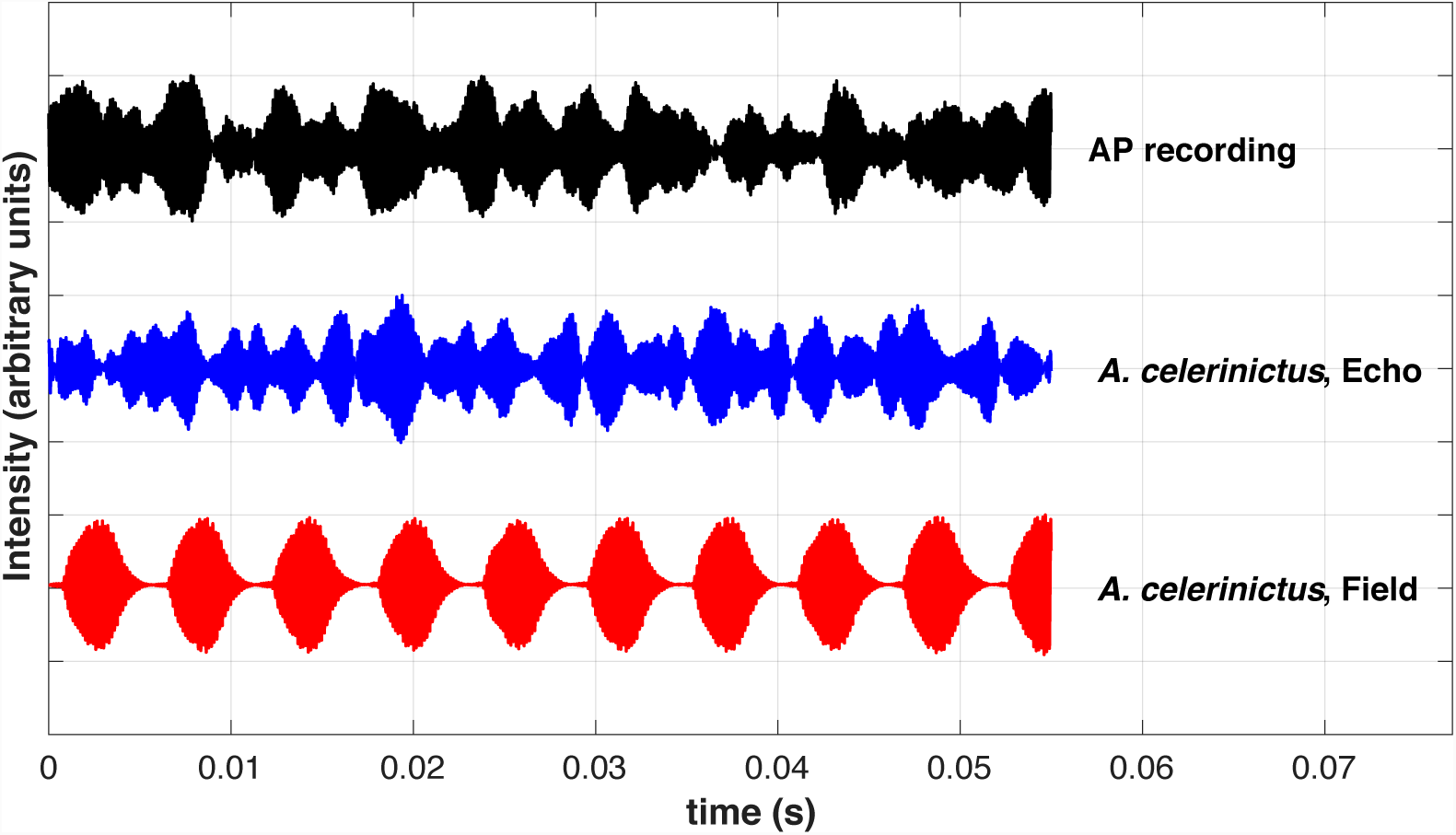
Pulse structure of AP recording and *A. celerinictus* with and without echo. This plot shows the pulse structure of the three recordings over time. The AP recording (**top, black**) exhibits an irregular pulse structure that does not match insect calls from isolated field recordings. The *A. celerinictus* call (**bottom, red**) is highly uniform in pulse structure when recorded in the field however this is modified by internal echoes when recorded indoors. The *A. celerinictus* call recorded indoors (**middle, blue**) is an excellent match to the AP recording in pulse structure, pulse repetition rate, pulse repetition rate stability, and amplitude spectrum.

The pulse structure labeled “*A. celerinictus,* Echo” in the figures results from a recording made in a house with tile floor and drywall construction. The pulse-envelope-structure of both the AP recording and the recordings of an *A. celerinictus* call played indoors are not constant through time, which can be due to complicated interference patterns that result from multiple sound pulses superimposed on one another with pulse-to-pulse variation in the phases of the interfering 7 kHz components. Extended Data Figure 1 shows a longer timescale of pulse structures, and Extended Data Figures 2 and 3 show quantitatively the similarity between the echoed *A. celerinictus* call and the AP recording. Extended Data Figure 4 shows a similar resulting pulse structure from an echoed recording of related *Anurogryllus muticus* obtained by A.L.S. in Costa Rica from within a restaurant compared with a field recording of the same species. These analyses all show that the pulse structure of the AP recording is consistent with an echoing cricket call. A.L.S. also notes that while crickets calling away from structures were fairly easy to locate, the complex sound environment and echoes made it very difficult to find individual crickets calling near buildings.

In *A. celerinictus* the file has between 40-50 teeth spread over 2.5-3.0 mm^20^, and the number of cycles in each pulse is around 30 as shown in Figure 5. The number of oscillations per pulse for the field recording of the insect matches that seen in the AP recording. This agreement is an independent additional piece of evidence for *A. celerinictus* being the source of the sound in the Cuba recording.

**Fig. 5.**
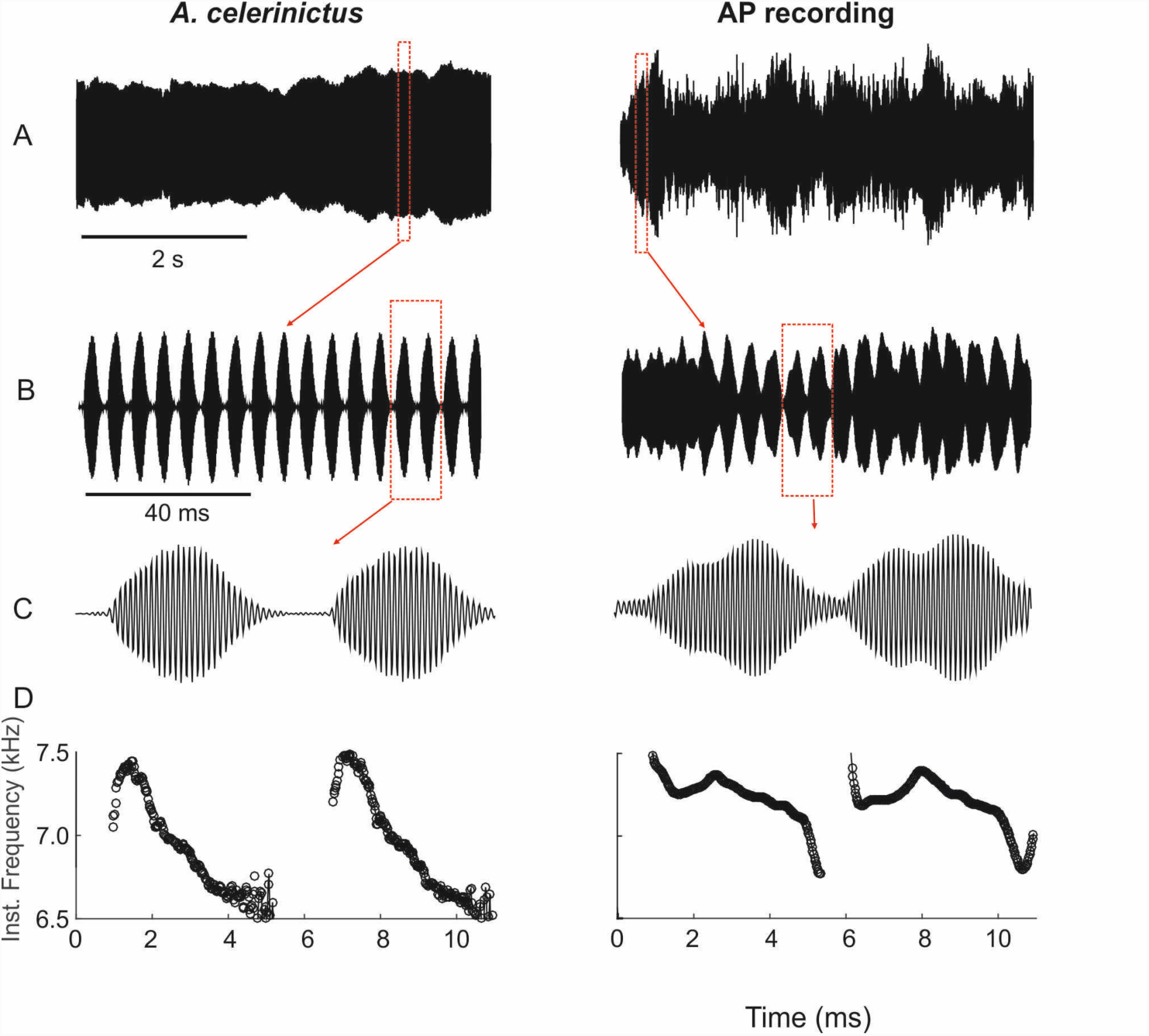
Frequency evolution and number of oscillations within a pulse. **A**, **B**, and **C**, show the time series with a scale bar of 2 seconds, 40 ms and 4 ms respectively, for field and AP recordings. Panel **D** shows the frequency decay through one pulse, measured via the interval between zero crossings. In both the *A. celerinictus* recording and AP recording the frequency decays over the sound pulse.

Figure 5 compares the decay of the instantaneous frequency over the course of a pulse. This is due, in part, to deceleration of the cricket’s wing through each pulse. While individual pulses in the AP recording are impacted by echoes of the preceding sound pulse, there is a clear decay in frequency in both cases.

The 7 kHz buzzing sound recorded by U.S. embassy personnel and released by the AP is entirely consistent with an echoing insect source, and not likely to have resulted from a “sonic attack.” Other hypotheses that invoke stable digital signal sources^13–16^ for this sound (a) do not explain the few-percent drift in the PRR, (b) are not as well-matched spectrally^14^, and (c) fail to explain the pulse structure and frequency decay through each pulse seen in the AP recording. The first individual to believe this sound was associated with health issues reported^1^ that the sound stopped abruptly when he opened the front door. This and other reports of the sound abruptly stopping with movement in a room^25^ are also consistent with an insect stopping a call when threatened.

The situation in Cuba has^1^ understandably led to concern and anxiety, and the sonic attack hypothesis has gained widespread attention in the media. However, this paper shows that sounds like those in the AP recording have a natural explanation. In particular, we have six lines of evidence to show that the sounds recorded by U.S. personnel in Cuba correspond to the calling song of a specific cricket, with echoes. The following quantitative signal characteristics provide independent lines of evidence to support the conclusion that the sound recorded by U.S. personnel in Cuba is of biological origin:

1. Carrier frequency of 7 kHz
2. Pulse repetition rate of 180 Hz
3. Timescale and amount of pulse repetition instability
4. Echo phenomenology
5. Number of oscillations per pulse
6. Frequency decay of about 1 kHz over pulse duration

Thus, while disconcerting, the mysterious sounds in Cuba are not physically dangerous and do not constitute a sonic attack. The fact that the sound on the recording was produced by a Caribbean cricket does not rule out the possibility that embassy personnel were victims of another form of attack. While the causes of any signs and symptoms affecting U.S. personnel in Cuba are beyond the scope of this paper, a biological origin of the recorded sounds motivates a rigorous examination of other possible origins, including psychogenic, of reported neuro-physiological effects. This episode has potential parallels with a previous incident in U.S. history, “yellow rain” in Southeast Asia, where alleged chemical attacks were later determined to be of benign biological origin. In that instance bees, rather than crickets, were to blame.

## Methods

### A: The search for a biological source consistent with the AP recording

A wide diversity of organisms use sound as a method of communication, and particularly in biodiverse areas like Cuba there are many potential natural acoustic sources. Since these incidents were predominantly reported^2^ at night, primarily diurnal sound sources were eliminated from consideration. There might be more than one organism capable of reproducing a sound with the properties of the AP recording. This section explores the rationale for the selection of biological sources that were subjected to additional spectral analysis and comparison with the AP sample.

### Vertebrates-

#### Frogs

The Caribbean region does have loud frogs such as the Puerto Rican common coqui, *Eleutherodactylus coqui* (Bello and Espinosa 1871), well known for being introduced in Hawaii and apparently depressing property values^26^ due to their loud advertisement call. Frogs of the genus *Eleutherodactylus* do not produce continuous calls, however, nor do any other amphibians that might be encountered in the Caribbean. The presence of a distinctive pulse repetition rate in the AP recording makes it unlikely a chorus of individual frogs (or other organisms) was responsible.

#### Birds

There are no continuously calling nocturnal birds. Additionally, most bird song is of high-complexity to aid in species recognition. Birds are not capable of producing a sustained noise for minutes.

### Insects-

The identification of potential insect sound sources was helped immensely by the website Singing Insects of North America (SINA) https://entnemdept.ifas.ufl.edu/walker/Buzz,maintained^22^ by Thomas J. Walker. Prof. Walker has also conducted extensive work in and published on the calling insects of Caribbean islands. Due to political considerations involving the complicated relationship between the United States and Cuba, there is comparatively little publicly-available data on calling insects of Cuba. This study focused on insects that are present on other Caribbean islands and in Florida in the hope that if a close match were found this would narrow the search for potential Cuban insect sources.

There are three primary groups of relevant calling insects: cicadas (superfamily Cicadoidea), katydids or bush crickets (family Tettigoniidae), and various kinds of crickets (superfamily Grylloidea). Many insects (e.g., cicadas, crickets, katydids) communicate acoustically, and exploit both audio and ultrasonic signals. Among these, crickets (one of the most studied models of acoustic communication) exhibit behavioural and biophysical aspects that have fascinated humans for decades.

#### Cicadas

The Cuban government has implicated^27^ cicadas as a potential source for the sound heard by U.S. personnel. Unfortunately, little is published about the songs of Cuban cicadas, making detailed spectral analysis difficult. Multiple press reports mention^2^ that the sounds reported by U.S. personnel were heard at night, and cicadas are largely diurnal callers, making it unlikely that they are responsible.

#### Katydids

*Neoconocephalus robustus* is the only katydid with a publicly-available recording and an appropriate pulse repetition rate and carrier frequency. Katydids found in the Caribbean typically have^21^ a wider power spectrum than “pure tone” producing crickets. As seen in Figures 1 and 3, *N. robustus* has a very different acoustic signature to the AP recording. The largest inconsistency is the lack of a “picket fence” series of lines in the amplitude spectrum. This is an indication of the instability in the PRR of the katydid call. While *N. robustus* is not known to be present on Cuba^28^ it seemed appropriate to include a katydid exemplar for analysis with a similar carrier frequency and pulse rate as the AP recording, as an illustration of what katydid calls looked like compared to crickets. There are multiple species of *Neoconocephalus* and *Conocephalus* katydids in Cuba. The lack of available audio recordings makes further analysis difficult, however an initial analysis suggests that instability in the PRR is characteristic of North American katydids with recordings uploaded to the SINA^22^ website.

#### Crickets

Male crickets produce three types of signals: 1) Calling songs attract distant females, and are loud, continuous pure-tone calls (with narrow frequency spectrum) peaking between 3-8 kHz, depending on the species^22^. 2) Courtship songs are whisper-like signals of low intensity, given at frequencies higher than calling songs (10-15 kHz). 3) Aggressive or rivalry signals are produced in the presence of other males, usually with broadband spectra^22^. The calling song informs the female of the male’s presence, its genetic compatibility, and location. Such information is conveyed in the loudness, clarity, directionality, and power spectrum of the call. The calling song’s dominant frequency is often within the human hearing range (50 Hz - 20 kHz), and is the call type that is relevant to this research. Crickets are a monophyletic group with multiple subfamilies. After combing through the species accounts of over 130 crickets on the SINA website^22^, only one species was encountered with a pulse repetition rate matching that of the AP recording. As noted by Walker in his 1973 description of the species, *A. celerinictus* has^20^ the fastest wing stroke rate known for a continuously calling cricket. This turned out to be the best match with the AP recording. Allard, with a delightful turn of phrase, describes^29^ encountering an *Anurogryllus* cricket in the Indies as follows:

“In the Dominican Republic when the warm and humid evening arrives, scattered chirping and tinkling notes issue from the shrubs and trees here and there. Some of these are clear, incisive little points of high-pitched sound; others are powerful, penetrating, buzzing, almost ringing noises, continuous and even very disconcerting to many people because of the incessant din.

In the Capital city, Ciudad Trujillo, the large brown cricket *Anurogryllus muticus* (DeGeer) is very common and noisy throughout the winter. As soon as the night came on and lights appeared, these ubiquitous crickets began their activities out-of-doors in the yard and even within the wide-open houses, for there are no screened windows or doors in the typical Spanish houses.

The song of the males of this cricket, here, is a continuous ringing z-z-z-z-z-z-of tremendous volume and penetration which practically fills a room with veritable din. The song is quite like that of our common cone-head, *Neoconocephalus robustus crepitans* (Scudder) of the eastern United States. After being accustomed to hear the trilling notes, definitely musical in tonality, of our American individuals of this species, I was somewhat nonplussed to hear this tropical cricket singing continuously, with all the characteristics of a cone-headed katydid, and with no tonality in its stridulation.”

### A note on genus *Anurogryllus* including the status of *A. celerinictus* in Cuba

Publically-available call data exists for only 2 crickets of genus *Anurogryllus* in the Caribbean: *Anurogryllus celerinictus* (Walker 1973) and *Anurogryllus muticus* (De Geer 1773). Prior to 1973 the species *A. muticus* was thought to range from the type locality in Suriname through the Eastern United States, but *A. celerinictus* and *A. arboreus* were described^20^ by Walker in 1973 when he split the complex into three species. The primary evidence for this was that crickets previously known as *A. muticus* produced calling songs with three distinctive wingstroke frequencies (or pulse repeat rates in the terminology used here). Walker recorded calls of *A. celerinictus* from Jamaica, Grand Cayman, and Big Pine Key. He found that these calls ranged in wingstrokes per second from ∼145 Hz at 20 C to ∼190 Hz at 28 C with peak carrier frequency ranging from 6-7.4 kHz. Walker reported that *A. muticus* has a peak carrier frequency of 5.8-7.2 Hz but only reached a maximum wing stroke rate (PRR) of ∼150 Hz at 27 C with a minimum of ∼110 Hz at 20 C. *A. arboreus* (Walker 1973) is found only in the mainland U.S. and has a wing stroke rate (PRR) of under 80 Hz. As detailed^20^ in Walker’s 1973 description, *A. celerinictus* and *muticus* are not easily distinguishable based on morphology, thus records of *Anurogryllus* crickets listed^28^ as the earlier described *A. muticus* from Cuba may well include individuals of *A. celerinictus.* As there is little published information regarding the calls of Cuban *Anurogryllus* species, it may well be that an individual of another species for which there is no publicly-available call data also produces a similar call. Given the information available and the presence of *A. celerinictus* in the Caymans, Florida Keys, and Jamaica, it seems reasonable to expect that populations previously referred to as *A. muticus* from Cuba might have the *A. celerinictus* call type and therefore be representatives of *A. celerinictus.* Indeed in Walker’s description of *A. celerinictus* he postulated that the specimens found in the Florida Keys might have recently emigrated from Cuba^20^ as subsequent trips to the same localities did not produce additional *A. celerinictus*. Analysis of the AP recording from Cuba with a higher wingstroke rate than *A. muticus* as reported by Walker (1973) provides evidence for the presence of *A. celerinictus* on the island of Cuba.

Field crickets (Gryllinae) of genus *Gryllu*s are well studied, particularly the Jamaican field cricket *Gryllus assimilis. G. assimilis* produces calls with a chirp rate^23,24^ of about once per second and most North American biologists (or members of the public) would immediately recognize this call as a cricket. It is unclear why *G. assimilis* was implicated by the Cuban report^12^ when the song is readily available via multiple sources online and sounds qualitatively different from the recording released by the AP. A quantitative analysis, shown in Fig. 2, reinforces this conclusion.

### Recordings used in the analysis

#### AP Recording

An.mp4 file was extracted from the AP’s posted^30^ recording https://www.youtube.com/watch?v=Nw5MLAu-kKs&feature=youtu.be using the program FonePaw Video Converter Ultimate version 2.25. The first 0.25 seconds of the AP recording was trimmed as there was no signal. Similarly, the end of the file without signal was trimmed using Audacity 2.2.2 to generate a final.wav file of 5.11 seconds duration. The AP recording was released as both a long format video with additional information^5^, and as a standalone.mp4 file^30^. In the long-format video the AP states that they received multiple similar recordings and that “the U.S. embassy in Havana has played^5^ these recordings for Americans who are working there so they know what to listen to.” The accompanying AP story^5^ asserts that these recordings were received from a U.S. government employee, and were sent to the U.S. Navy for acoustic analysis. The Cuban analysis^12, 27^ shows a coarse power spectrum with a 7 kHz peak. This supports the conclusion (in agreement with references^14–16^) that the AP released recording is representative of the “sonic attack” recordings in Cuba.

#### *A. celerinictus* field recording

The recording of *A. celerinictus* was downloaded from the SINA website at http://entnemdept.ufl.edu/Walker/buzz/492a.htm. This file is 20 seconds of calling song recorded by T. J. Walker from Big Pine Key, Monroe County, FL. The temperature was 27 C at the time of recording. This file is referred to as “*A. celerinictus* field” in the figures and manuscript.

#### *A. celerinictus* echo recording

This recording was generated by playing the “*A. celerinictus* field” recording on a UE Wonderboom speaker at the base of a stairwell in a house with drywall walls and a tile floor. Other locations indoors in the same tile floored house produce similar results. A recording was also made outdoors to verify the speaker did not introduce distortions or false echoes.

#### Neoconocephalus robustus

This recording was also downloaded from the website of T. J. Walker at URL http://entnemdept.ufl.edu/walker/buzz/195a.htm and is a male from Washington County Ohio calling at 23.8 C.

#### Anurogryllus muticus

A.L.S. made a series of recordings of *A. muticus* in the Pacific Coast lowlands of Costa Rica in December 2018. Two recordings are presented here as representative of an *Anurogryllus* species recorded away from human structures, Extended Data Audio 2, and from within an open-air restaurant, Extended Data Audio 3. These recordings are illustrative as they show in Extended Data Figure 4 how an echoed recording of an *Anurogryllus* cricket has a obscured pulse structure in comparison to a field recording made away from buildings. A.L.S. also notes that calling *A. muticus* in Costa Rica is loud enough to be the dominant sound even when calling outside noisy restaurants. The only major difference between the call of *A. muticus* and *A. celerinictus* is the higher pulse repetition rate of *A. celerinictus.* To A.L.S. the two calls sound similar, and if both species are on Cuba as they are on Jamaica^20^ it is possible that U.S. personnel may have heard both species. A release of any additional recordings made by U.S. personnel would clarify this point.

## Supporting information

ExtendedDataAudio2

ExtendedDataAudio3

ExtendedDataAudio1

## Analysis methodology

### Making Figure 1

The four audio files described above were each loaded into MATLAB and trimmed to the first 5 seconds to match the length of the AP recording. As these audio files were sampled at 44,100 samples/second, each 5 second clip has 220,500 data points. The average value of each dataset was subtracted prior to running the MATLAB FFT to obtain an amplitude spectrum, with no window function applied. This produces a spectral bin size of 0.2 Hz. The amplitude spectra were averaged over 25 adjacent data points, to suppress fluctuations, giving an effective spectral resolution of 5 Hz. Each smoothed spectrum was normalized to its peak value. MATLAB program (to be posted online pending publication) Cuba_Figure1.m was used to create the figure.

### Making Figure 2

Peak emission frequencies and PRR values were measured either using Audacity version 2.2, or with custom MATLAB code, on downloaded audio files as listed in Supplementary Table 1. Relevant MATLAB programs will be posted online pending publication.

### Justification for species and associated data plotted in Figure 2

The recordings available at the SINA website^22^ were primarily used to determine PRR and peak emission frequency for species of interest. The provenance and results are shown in Extended Data Table 1.

### Making Figure 3

Subsets of 8,192 data points, compromising 0.1858 seconds, of each 5-second recording were sequentially analyzed. The MATLAB FFT function was used to create 5.38 Hz wide spectral bins. Then the MATLAB script Cuba_Figures.m incremented by 4096 data points (half of one sub-segment length) to compute the next FFT, iterating through the entire data file. This means that there is intentional overlap between adjacent subsamples, as a way to smooth the data for visualization. The first of these subspectra was plotted at the top of each waterfall plot. This was scaled to the peak amplitude in the range between 5.5 and 8.5 kHz. The MATLAB program (to be posted online pending publication) Cuba_Figures.m was used to create the figure.

### Making Figure 4

This figure was created by simply plotting intensity vs. time for each of the three data sets using the same MATLAB program Cuba_Figures.m. This plot starts at 4 seconds into each recording.

### Making Figure 5

Panels A, B, C are simply different timescale representations of the two recordings. Panel D shows the instantaneous frequency of the signals, measured as Zero Crossing, was obtained with Hilbert transform, using the following MATLAB custom code.

### hilbert_transform.m

function[env, phase, inst_freq]=hilbert_transform(signal, Fs)

%Fs = sampling frequency in Hz

signal=signal/max(abs(signal));

signal=signal-mean(signal);

hilb=hilbert(signal);

env=abs(hilb); phase=atan2(imag(hilb),real(hilb)) + (pi/2);

inst_freq=(diff(unwrap(phase))./(1/Fs))./(2*pi);

phase=phase*180/pi;

phase=modrange(phase, −180, 180);

end

### C. Echo detection

If the pulse repetition rate and carrier phase were perfectly stable and the microphone and source were stationary, one would expect echoes to arrive after a constant delay following each pulse. Given the pulse structure evolution seen in Extended Data Figure 1 in the AP sample and the *A. celerinictus “*Echo” recording, it appears this is not the case, even for the interior experiment where the source and microphone were stationary.

Pulse-to-pulse variability evidently induces constructive and destructive interference patterns that vary over time. Adding the field recording to a time-delayed replica reproduces the echo phenomenology. This was verified using Audacity 2.2. With this qualitative understanding of the effect, a quantitative assessment was undertaken. The first step, shown in Extended Data Figure 2, was to identify peaks in the recorded sound intensity. The interval between successive peaks was measured using the MATLAB function findpeaks() in the script EchoInterval.m (available upon publication). Peak detection criteria were:

- Minimum separation between peaks of 0.5 ms.
- Minimum peak height of normalized power 0.12.
- Minimum peak width 0.5 ms.

As shown in Extended Data Figure 3 the distribution of peak-to-peak intervals is an indicator of echoes and interference. A “short” peak-to-peak time indicates that an echo interrupts the original pulse train. A “long” peak-to-peak time indicates that destructive interference suppresses a peak. In both the AP and *A. celerinictus* “Echo” recordings the fraction of “short” and “long” intervals is similar. This supports the conclusion that the AP recording arises from echoes of a natural source.

## Extended Data Table

**Extended Data Table 1.**
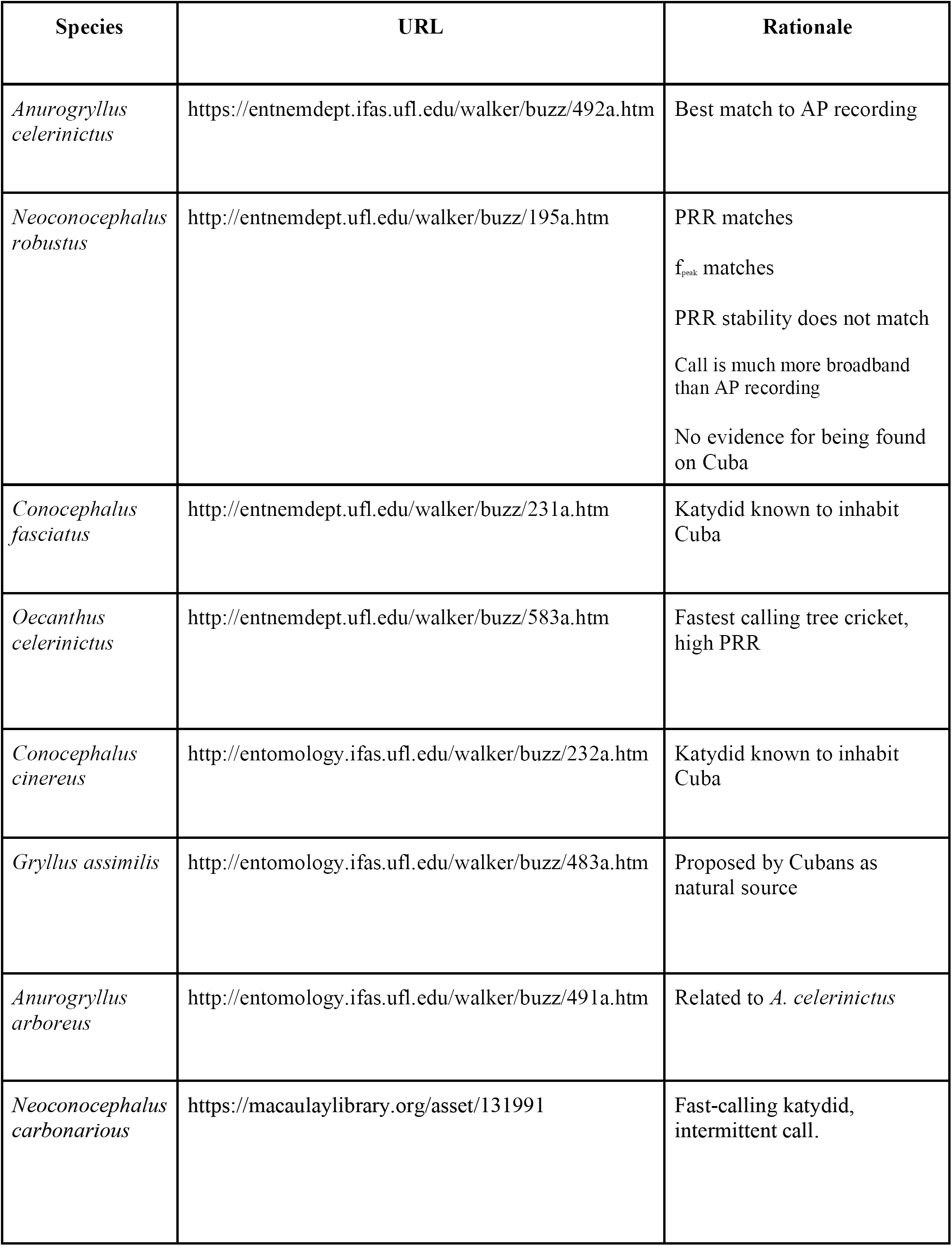
Full genus and species of all data points in Fig. 2 along with justification for inclusion.

## Extended Data Figures

**Extended Data Figure 1.**
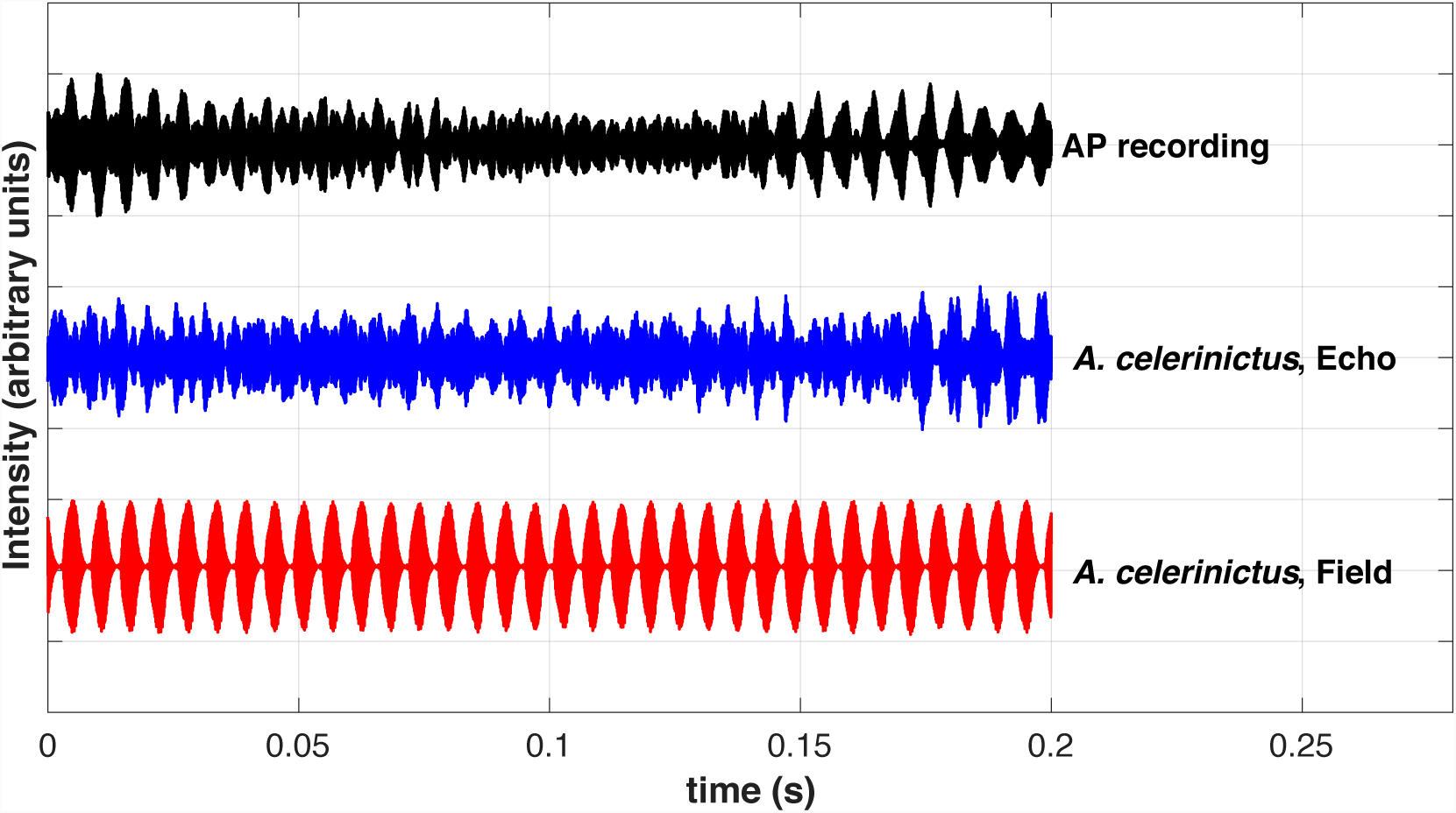
Expanded pulse structure comparison. This shows a longer time series for all three recordings of interest. The red *A. celerinictus* Field recording converts into the blue Echo trace when played indoors, which strongly resembles the AP recording. In the AP recording there are regions of high pulse amplitude at 0.01 and 0.17 seconds in this plot bounding a region of low pulse amplitude at 0.11 seconds. This seemingly symmetrical pattern suggested interfering waves moving in and out of phase.

**Extended Data Figure 2.**
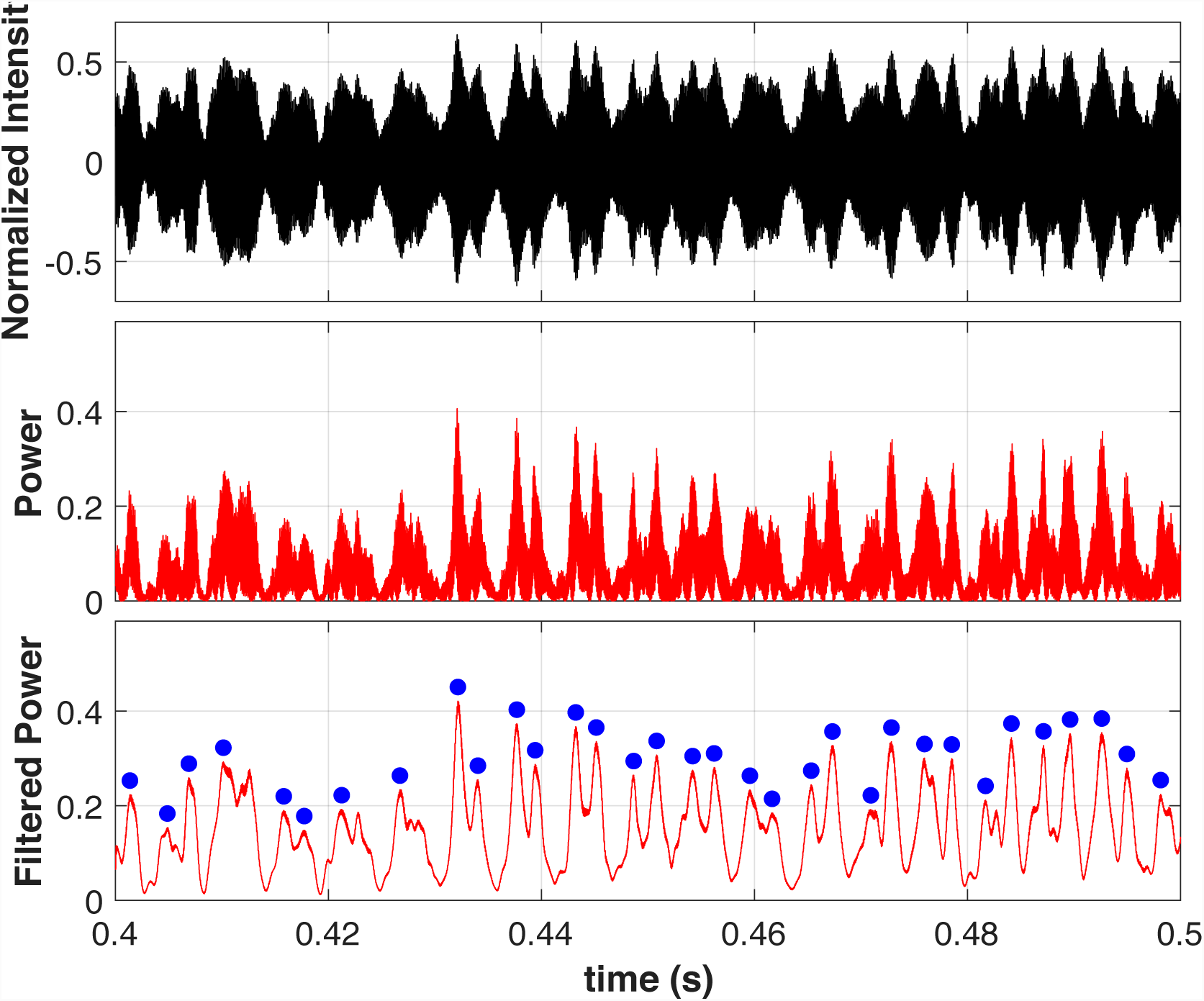
Pulse interval determination. The upper panel shows intensity vs. time from the AP recording, the middle panel shows power (intensity squared), the lower panel shows a low-pass filtered version of the power vs. time. This allows for automated peak detection (shown as blue dots), from which the interval between peaks can be determined.

**Extended Data Figure 3.**
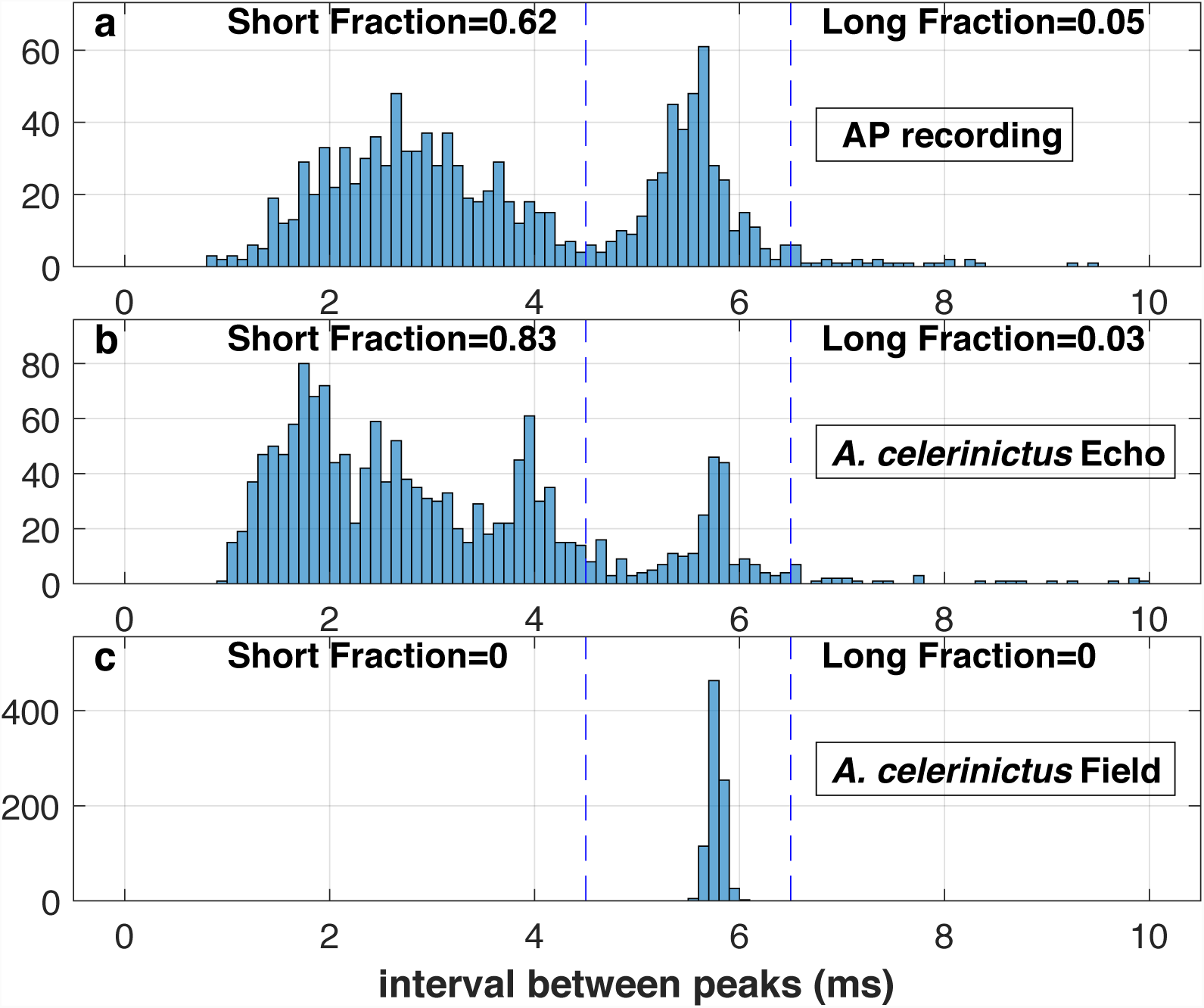
Peak interval histograms. **c**, In the Field recording the interval between two successive peaks is constant at just under 6 ms, the reciprocal of the PRR. **b**, A significant fraction (0.83) of “interloper” peaks from the echo produce short intervals. Also, ∼3% of the time destructive interference suppresses adjacent peaks. **a**, Similar interval statistics occur in the AP recording. **a-c**, The vertical dashed lines define short and long fractions respectively.

**Extended Data Figure 4.**
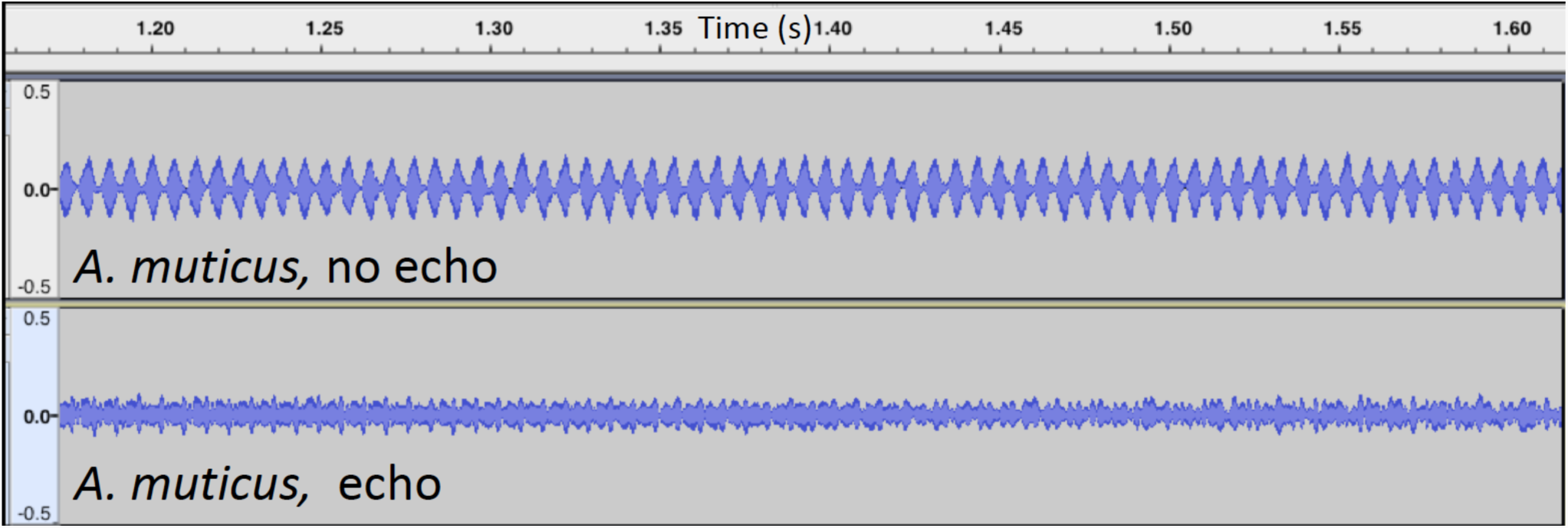
Recording of related *Anurogryllus muticus* from Costa Rica with and without echoes. The top panel shows a recording of *A. muticus* taken in a palm grove away from structures that might produce echoes. The bottom panel is a recording of the same species made approximately 2 meters away from a concrete wall inside a restaurant in Costa Rica. This shows a real-world example of how recordings near structure obscure the pulse structure of *Anurogryllus* calls.

**Extended Data Audio 1. Echoed recording of *A. celerinictus*.** This 5 second audio clip was obtained by replaying the field recording indoors.

**Extended Data Audio 2. Field recording of *Anurogryllus muticus* from Costa Rica without echoes.** This recording was made in a stand of coconut palms away from any buildings and therefore is without echoes. This pulse structure looks very similar to the *A. celerinictus* “field” recording in pulse structure, though the pulse repetition rate is slower.

**Extended Data Audio 3. Field recording of *A. muticus* from Costa Rica made near a cement wall.** This recording was made inside a restaurant in Costa Rica, where a cricket was calling just outside adjacent a cement wall. Here there are considerable echoes that impact the pulse structure, in much the same way as the AP recording.

## Supplementary information line

Extended Data Table 1

Extended Data Figs. 1-4

Extended Data Audio 1-3

Caption for Extended Data Audio 1-3

## Acknowledgements

T. J. Walker has devoted his career to entomological scholarship and public accessibility of insect call recordings. The NSF GRFP and the UC Berkeley Museum of Vertebrate Zoology provided support to A.L.S. who also thanks Jimmy McGuire for his mentorship and the many opportunities for field expeditions listening to frogs and crickets. David Wake and Alexander Loomis provided extensive comments and advice on this manuscript. FMZ is funded by the European Research Council, ERC-CoG-2017-773067.

## Competing interests

The authors declare no competing interests.

Correspondence to Alexander Stubbs at

## Data Disposition

All recordings used for analysis except for the the three supplementary audio recordings are publicly available. The “*A. celerinictus* echo” recording is available as “Extended Data Audio 1 echoed_celerinictus.wav” while the *A. muticus* echo and *A. muticus* field are available as Extendeddataaudio1.mpg and Extendeddataaudio2.

All code used for spectral analysis and to make the figures will be uploaded to github upon publication. The data used to generate Figure 2 is part of this code.

